# Metaepigenomic analysis reveals the unexplored diversity of DNA methylation in an environmental prokaryotic community

**DOI:** 10.1101/380360

**Authors:** Satoshi Hiraoka, Yusuke Okazaki, Mizue Anda, Atsushi Toyoda, Shin-ichi Nakano, Wataru Iwasaki

## Abstract

DNA methylation plays important roles in prokaryotes, such as in defense mechanisms against phage infection, and the corresponding genomic landscapes—prokaryotic epigenomes—have recently begun to be disclosed. However, our knowledge of prokaryote methylation systems has been severely limited to those of culturable prokaryotes, whereas environmental communities are in fact dominated by uncultured members that must harbor much more diverse DNA methyltransferases. Here, using single-molecule real-time and circular consensus sequencing techniques, we revealed the ‘metaepigenomes’ of an environmental prokaryotic community in the largest lake in Japan, Lake Biwa. A total of 19 draft genomes from phylogenetically diverse groups, most of which are yet to be cultured, were successfully reconstructed. The analysis of DNA chemical modifications identified 29 methylated motifs in those genomes, among which 14 motifs were novel.

Furthermore, we searched for the methyltransferase genes responsible for the methylation of the detected novel motifs and confirmed their catalytic specificities via transformation experiments involving artificially synthesized genes. Finally, we found that genomes without DNA methylation tended to exhibit higher phage infection levels than those with methylation. In summary, this study proves that metaepigenomics is a powerful approach for revealing the vast unexplored variety of prokaryotic DNA methylation systems in nature.

## Introduction

DNA methylation is a major class of epigenetic modification that is found in diverse prokaryotes, in addition to eukaryotes^1^. For example, prokaryotic DNA methylation by sequence-specific restriction-modification (RM) systems that protect host cells from invasion by phages or extracellular DNA has been well characterized and is utilized as a key tool in biotechnology^2,3,4^. In addition, recent studies have revealed that prokaryotic DNA methylation plays additional roles, performing various biological functions, including regulation of gene expression, mismatch DNA repair, and cell cycle functions^5–9^. Research interest in the diversity of prokaryotic methylation systems is therefore growing due to their importance in microbial physiology, genetics, evolution, and disease pathogenicity^7,10^. However, our knowledge of the diversity of prokaryotic methylation systems has been severely limited thus far because most studies must focus only on the rare prokaryotes that are cultivable in laboratories.

The recent development of single-molecule real-time (SMRT) sequencing technology provides us with another tool for observing DNA methylation. An array of DNA methylomes of cultivable prokaryotic strains, including N6-methyladenine (m6A), 5-methylcytosine (m5C), and N4-methylcytosine (m4C) modifications, have been revealed by this technology^11–14^. Despite its high rates of base-calling and methylation-detection errors per raw read^15,16^, SMRT sequencing technology can produce ultralong reads of up to 60 kb with few context-specific biases (*e.g.*, GC bias)^17^. This characteristic enables SMRT sequencing to achieve high accuracy by merging data from many erroneous raw reads originating from clonal DNA molecules, typically from cultivated prokaryotic populations^18^. Alternatively, in an approach referred to as circular consensus sequencing (CCS), a circular DNA library is prepared as a sequence template to allow the generation of a single ultralong raw read containing multiple sequences (‘subreads’) that correspond to the same stretch on the template^19,20^; therefore, a cultivated clonal population is not required to achieve high accuracy^21^. However, CCS has thus far been applied in only a few shotgun metagenomics studies^22^ and, to the best of our knowledge, has not yet been applied to ‘metaepigenomics’ or direct methylome analysis of environmental microbial communities, which are usually constituted by uncultured prokaryotes.

Here, we applied CCS to shotgun metagenomic and metaepigenomic analyses of freshwater microbial communities in Lake Biwa, the largest lake in Japan, to reveal the genomic and epigenomic characteristics of the environmental microbial communities using the PacBio Sequel platform. Freshwater habitats are rich in phage-prokaryote interactions^23–26^, which are known to be closely related to prokaryotic DNA methylation. CCS analyses of the environmental microbial samples allowed reconstruction of draft genomes and the identification of their methylated motifs, at least 14 of which were novel. Furthermore, we computationally predicted and experimentally confirmed four methyltransferases (MTases) responsible for the detected methylated motifs. Importantly, two of the four MTases were revealed to recognize novel motif sequences.

## Materials and methods

### Sample collection

Water samples were collected at a pelagic site (35°13′09.5″N 135°59′44.7″E) in Lake Biwa, Japan (Fig. S1a) on December 26, 2016. The sampling site was located approximately 3 km from the nearest shore and had a depth of 73 m. The lake has a permanently oxygenated hypolimnion and was thermally stratified during sampling (Fig. S1b). Water sampling into prewashed 5-L Niskin bottles was conducted at depths of 5 m and 65 m, above and below the thermally stratified layer, respectively. The vertical profiles of temperature, dissolved oxygen concentrations, and chlorophyll *a* concentrations were measured using a conductivity, temperature, and depth probe *in situ*. Equipment that could come into direct contact with the water samples in the following steps was either sterilized by autoclaving or disinfected with a hypochlorous acid solution. The water samples were transferred to sterile bottles, kept cool in the dark, and immediately transported to the laboratory. Water samples with a total volume of approximately 30 L were prefiltered through 5-μm membrane PC filters (Whatman). Microbial cells were collected using 0.22-μm Sterivex filters (Millipore) and immediately stored at −20°C in a refrigerator until analysis.

### DNA extraction and SMRT sequencing

The microbial DNA captured on the Sterivex filters was retrieved using a PowerSoil DNA Isolation Kit (QIAGEN) according to the supplier’s protocol with slight modifications. The filters were removed from the container, cut into 3-mm fragments, and directly suspended in the extraction solution from the kit for cell lysis. The bead-beating time was extended to 20 minutes to yield sufficient quantities of DNA for SMRT sequencing, with reference to Albertsen *et al.*^27^ SMRT sequencing was conducted using a PacBio Sequel system (Pacific Biosciences) in two independent runs according to the manufacturer’s standard protocols. SMRT libraries for CCS were prepared with a 4-kb insertion length, and two SMRT cells were used for each sample as technical replicates.

### Bioinformatic analysis of CCS reads

Reads that contained at least three full-pass subreads were retained to generate consensus sequences (CCS reads) using the standard PacBio SMRT software package with the default settings. Only CCS reads with >97% average base-call accuracy were retained. For taxonomic assignment of the CCS reads, Kaiju^28^ in *Greedy-5* mode with the NCBI NR database^29^ and Kraken^30^ with the default parameters and complete prokaryotic genomes from RefSeq^31^ were used. CCS reads that potentially encoded 16S ribosomal RNA (rRNA) genes were extracted using SortMeRNA^32^ with the default settings, and the 16S rRNA sequences were predicted by RNAmmer^33^ with the default settings. The 16S rRNA sequences were taxonomically assigned using BLASTN^34^ searches against the SILVA database release 128^35^, where the top-hit sequences with e-values ≤1E-15 were retrieved.

CCS reads were *de novo* assembled using Canu^18^ with the *-pacbio-corrected* setting and Mira^36^ with the settings for PacBio CCS reads, according to the provided instructions. After removal of the assembled contigs that were suggested to contain repeats, the contigs were binned into genomes using MetaBAT^37^ based on genome coverage and tetra-nucleotide frequencies as genomic signatures, where the genome coverage was calculated by mapping the CCS reads to the binned genomes using BLASR^38^ with the settings for PacBio CCS reads. The quality of all genomes was assessed using CheckM^39^, which estimates completeness and contaminations based on taxonomic collocation of prokaryotic marker genes with the default settings. Sequence extraction and taxonomic assignment of 16S rRNA genes in each genome bin were conducted using RNAmmer^33^ with the default settings. Taxonomic assignment of the genome bins was based on the 16S rRNA genes if found or on the taxonomic groups most frequently estimated by CAT^40^ otherwise (and Kaiju^28^ if CAT did not provide an estimation).

Coding sequences (CDSs) in each genome bin were predicted using Prodigal^41^ with the default settings. Functional annotations were achieved through GHOSTZ^42^ searches against the eggNOG^43^ and Swiss-Prot^44^ databases, with a cut-off e-value ≤1E-5, and HMMER^45^ searches against the Pfam database^46^, with a cut-off e-value ≤1E-5. A maximum-likelihood (ML) tree of the genome bins was constructed on the basis of the set of 400 conserved prokaryotic marker genes using PhyloPhlAn^47^ with the default settings. Prophages were predicted using PHASTER^48^ with the default settings, and their sequence alignment was conducted using LAST^49^ with the default settings. CRISPR arrays were predicted using the CRISPR Recognition Tool^50^ with the default settings, and *cas* genes were annotated by querying 101 known CRISPR-associated genes in TIGRFAM^51^ using HMMER^45^ with a threshold of e-value ≤1E-5.

### Metaepigenomic and RM system analyses

DNA methylation detection and motif analysis were performed according to BaseMod (https://github.com/ben-lerch/BaseMod-3.0). Briefly, the subreads were mapped to the assembled contigs using BLASR,^38^ and interpulse duration ratios were calculated. Candidate motifs with scores higher than the default threshold value were retrieved as methylated motifs. Those with infrequent occurrences (<50) or very low methylation fractions (<1%) in each genome bin were excluded from further analysis.

Genes encoding MTases, restriction endonucleases (REases), and DNA sequence-recognition proteins were detected by BLASTP^34^ searches against an experimentally confirmed gold-standard dataset from the Restriction Enzyme Database (REBASE)^52^, with a cut-off e-value of ≤ 1E-15. Sequence specificity information for each hit MTase gene was also retrieved from REBASE.

### Experimental verification of MTase activities

Four estimated MTase genes (EMGBS3_12600, EMGBS15_03820, EMGBS10_10070, and EMGBD2_08790) were artificially synthesized with codon optimization and cloned into the pUC57 cloning vector by Genewiz (Table S1). The genes were subcloned into the pCold III expression vector (Takara Bio) using an In-FusionHD Cloning Kit (Takara Bio). The gene-specific oligonucleotide primers used for polymerase chain reaction and recombination are described in Table S2. For verification of the EMGBS10_10070 gene function, the 5’-ACGAGTC-3’ sequence was inserted downstream of the termination codon for the sake of the methylation assay (the first five-base ACG<u>**A**</u>G sequence was the estimated methylated motif, and the last five-base GAGTC is recognized by the restriction enzyme PleI) (Table S1).

The constructs were transformed into *Escherichia coli* HST04 *dam*^−^/*dcm*^−^ (Takara Bio), which lacks endogenous MTases. The *E. coli* strains were cultured in LB broth medium supplemented with ampicillin.

MTase expression was induced according to the supplier’s protocol. Plasmid DNAs were isolated using the FastGene Xpress Plasmid PLUS Kit (Nippon Genetics). SalI was employed to linearize the plasmid DNAs encoding EMGBS3_12600 and EMGBS15_03820 and then inactivated by heat. Methylation statuses were assayed by enzymatic digestion using the following restriction enzymes: BceAI and TseI for EMGBS3_12600, DpnII and XmnI for EMGBS15_03820, PleI for EMGBS10_10070, and FokI for EMGBD2_08790. All restriction enzymes were purchased from New England BioLabs. All digestion reactions were performed at 37°C for 1 h, except for those involving TseI (8 h) and FokI (20 min). Notably, although TseI digestion is conducted at 65°C in the manufacturer’s protocol, we adopted a temperature of 37°C to avoid cleavage of methylated DNA.

We further verified the methylated motifs that were newly estimated in this study, *i.e.*, those of EMGBS10_10070 and EMGBD2_08790. Chromosomal DNA was extracted from cultures of the transformed *E. coli* strains using a PowerSoil DNA Isolation Kit (QIAGEN) according to the supplier’s protocol. SMRT sequencing was conducted using PacBio RSII (Pacific Biosciences), and methylated motifs were detected via the same method described above.

### Data deposition

The raw sequencing data and assembled genomes were deposited in the DDBJ Sequence Read Archive and DDBJ/ENA/GenBank, respectively (Table S3). All data were registered under BioProject ID PRJDB6656.

## Results and discussion

### Water sampling, SMRT sequencing, and circular consensus analysis

Water samples were collected at a pelagic site in Lake Biwa, Japan, at 5 m (biwa_5m) and 65 m depths (biwa_65m), from which PacBio Sequel produced a total of 2.6 million (9.6 Gbp) and 2.0 million (6.4 Gbp) subreads, respectively (Table 1). The circular consensus analysis produced 168,599 and 117,802 CCS reads, with lengths of 4,474 ± 931 and 4,394 ± 587 bp, respectively (Table 1 and Fig. S2). In the shallow sample data, at least 90% of the CCS reads showed high quality (Phred quality scores >20) at each base position, except for the 5′-terminal five bases and 3′-terminal bases after the 5,638th base. In the deep sample data, the same was true, except for the 5′-terminal four bases and 3′-terminal bases after the 5,356th base (Fig. S3).

**Table 1.** Statistics of SMRT sequencing and CCS-read analysis.

| Sample | biwa_5m | biwa_65m |
| --- | --- | --- |
| Sequenced reads | 850,494 | 688,436 |
| Total base pairs (bp) | 9,570,723,004 | 6,419,717,083 |
| CCS reads | 168,599 | 117,802 |
| Read length (bp) | 4,474 ± 931 | 4,394 ± 587 |
| Total base (bp) | 754,416,328 | 517,663,806 |
| 16S rRNA | 170 | 106 |
| Length (bp) | 1,491 ± 64 | 1,468 ± 104 |

### Taxonomic analysis

Taxonomic assignment of the CCS reads was performed using Kaiju^28^ and the NCBI NR database^29^ (Fig. 1). The assignment ratios were >88% and >56% at the phylum and genus levels, respectively, which were higher than those for the Illumina-based shotgun metagenomic analysis of lake freshwater and other environments using the same computational method^28^. Kraken^30^ with complete prokaryotic and viral genomes in RefSeq^31^ (Fig. S4a-c) provided similar results but resulted in much lower assignment ratios (30% and 27%, respectively), likely due to the lack of genomic data for freshwater microbes in RefSeq. 16S rRNA sequence-based taxonomic assignment via BLASTN searches against the SILVA database^53^ also provided consistent results (Fig. S4d-f). It should be noted that 16S rRNA-based and CDS-based taxonomic assignments can be affected by 16S rRNA gene copy numbers and genome sizes, respectively.

**Figure 1.**
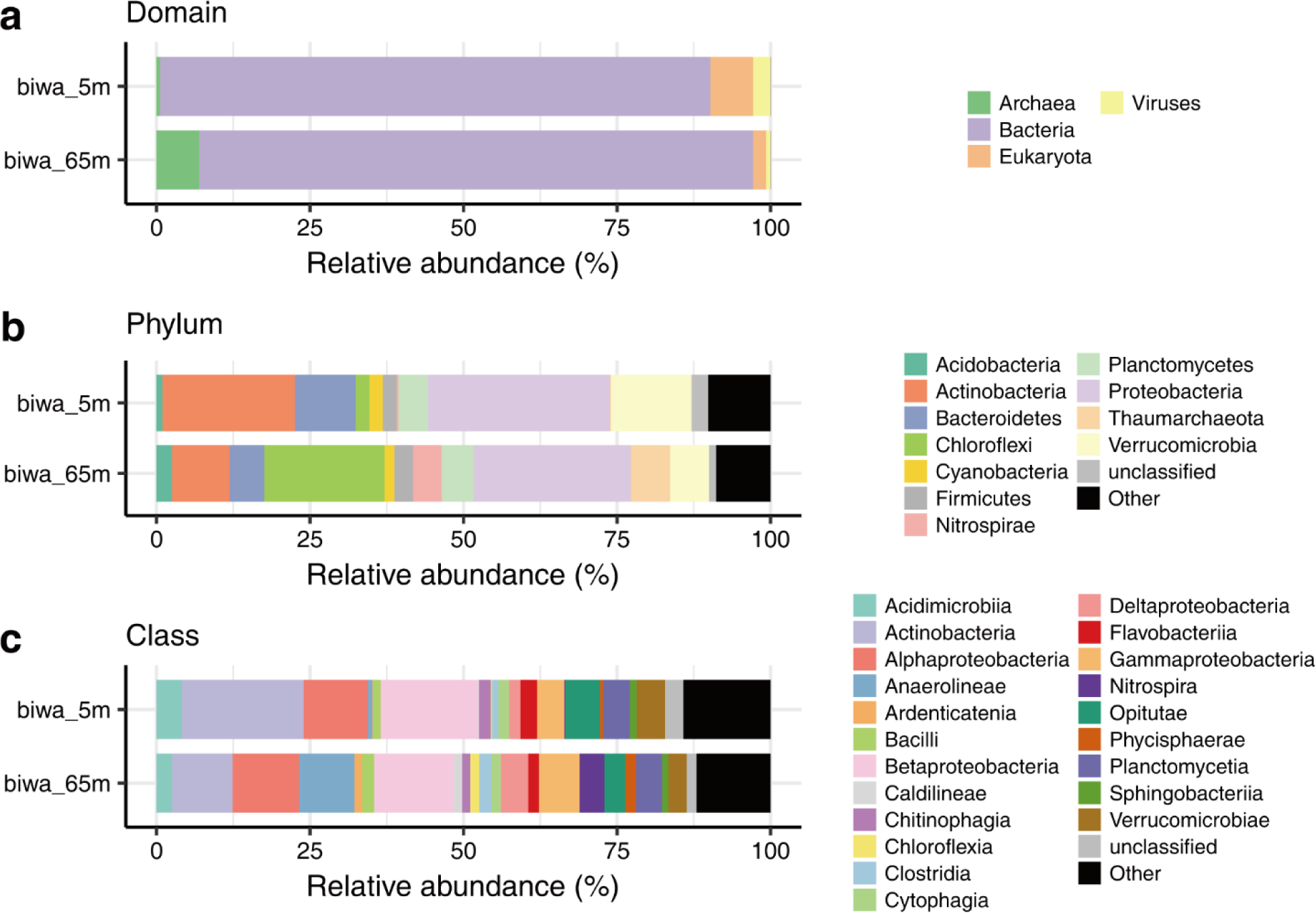
Phylogenetic distribution of CCS reads. Estimated relative abundances at the (**a**) domain, (**b**) phylum, and (**c**) class levels are shown. Eukaryotic and viral reads are ignored, and groups with <1% abundance are grouped as ‘Others’ in **b** and **c**.

At the phylum level, Proteobacteria dominated both samples, followed by Actinobacteria, Verrucomicrobia, and Bacteroidetes (Fig. 1). Chloroflexi and Thaumarchaeota were especially abundant in the deep water sample, consistent with previous findings^54,55^. The ratio of Archaea was particularly low in the shallow sample (0.6 and 6.9% in biwa_5m and biwa_65m, respectively). Although the filter pore-size range (5–0.2 μm) was not suitable for most viruses and eukaryotic cells, non-negligible ratios corresponding to their existence were observed in the shallow sample. The dominant eukaryotic phylum was Opisthokonta (2.68 and 0.92%), followed by Alveolata (1.67 and 0.45%) and Stramenopiles (1.45 and 0.15%). Among viruses, Caudovirales and Phycodnaviridae were the most abundant families in both samples. Caudovirales are known to act as bacteriophages, while Phycodnaviridae primarily infect eukaryotic algae. The third most abundant viral family was Mimiviridae, whose members are also known as ‘Megavirales’ due to their large genome size (0.6–1.3 Mbp)^56,57^. Viruses without double-stranded DNA (*i.e.*, single-stranded DNA and RNA viruses) were not observed because of the experimental method employed. Overall, the taxonomic composition was consistent with those obtained in previous studies on microbial communities in freshwater lake environments, reflecting the fact that SMRT sequencing provides taxonomic compositions consistent with those obtained using short-read technologies, such as the Illumina MiSeq and HiSeq platforms^58,59^.

### Metagenomic assembly and genome binning

The CCS reads from the shallow and deep samples were assembled into 554 and 345 contigs, respectively, using Canu^18^ (Table S4). The corresponding N50 values were 83 and 76 kbp, and the longest contigs had lengths of 481 and 740 kbp, respectively. Notably, the contigs were much longer than those obtained in a previous study that applied CCS for shotgun metagenomics analysis of an active sludge microbial community^22^. We also used Mira^36^ for metagenomic assembly, but this resulted in shorter longest contigs (148 and 151 kbp, respectively) and N50 values (19 and 18 kbp, respectively).

The contigs were binned to genomes using MetaBAT^37^, which is a reference-independent binning tool, based on CCS-read coverage and tetranucleotide frequency (Fig. 2 and Table 2). Among a total of 899 contigs, 390 (43.3%) were assigned to fifteen and four bins from the shallow and deep samples, respectively. We obtained a draft genome for each bin, where the completeness of the genome ranged from 17–99% (67% on average). Estimated contamination levels were low (<3% in each bin). Based on the total contig size and estimated genome completeness of each bin, the genome sizes were estimated to range from 1.0–5.6 Mbp. The GC content ranged from 29–68%, and the average N50 was 24 kbp, with a maximum of 1.67 Mbp.

**Table 2.**
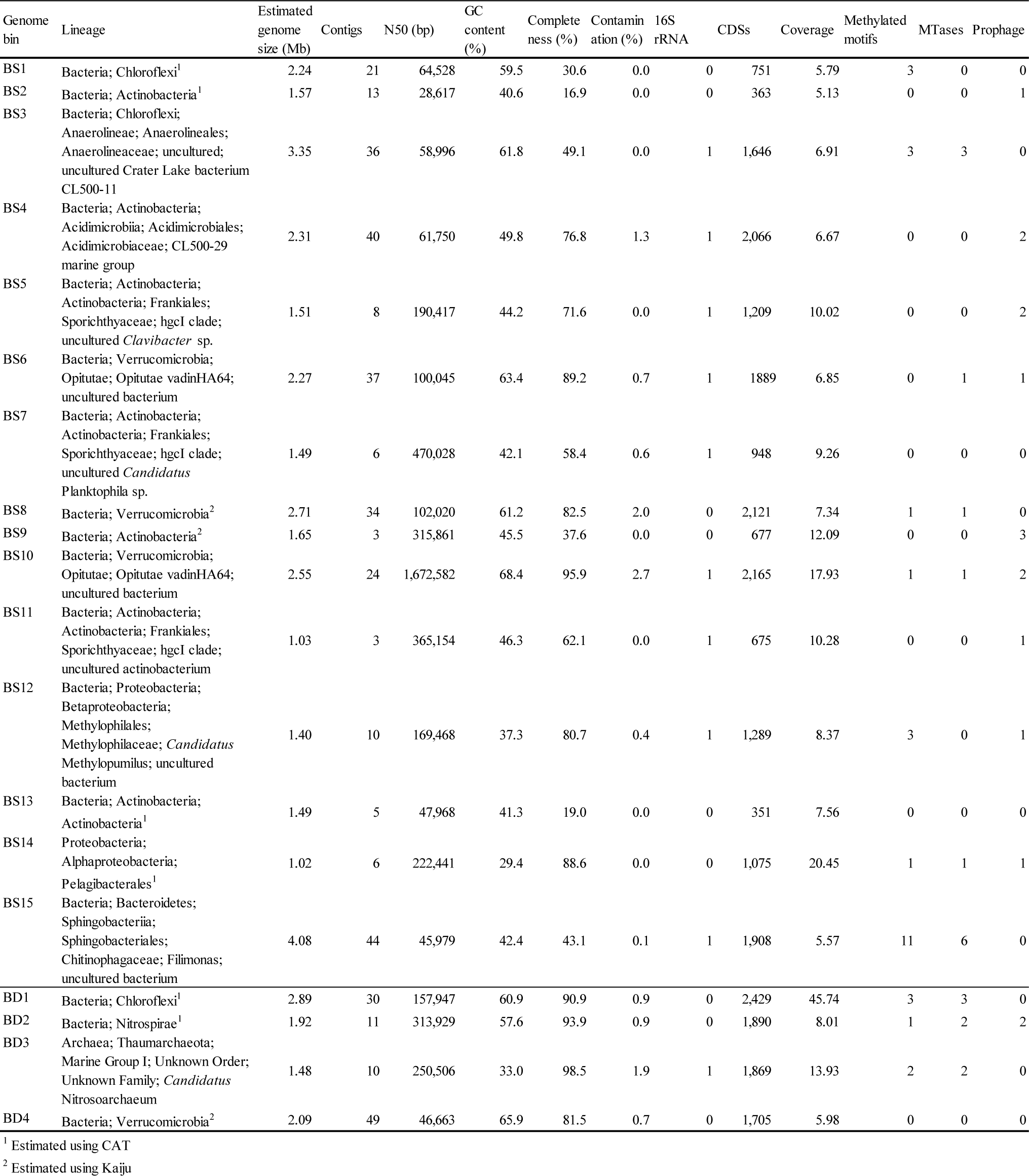
Statistics for genome bins.

**Figure 2.**
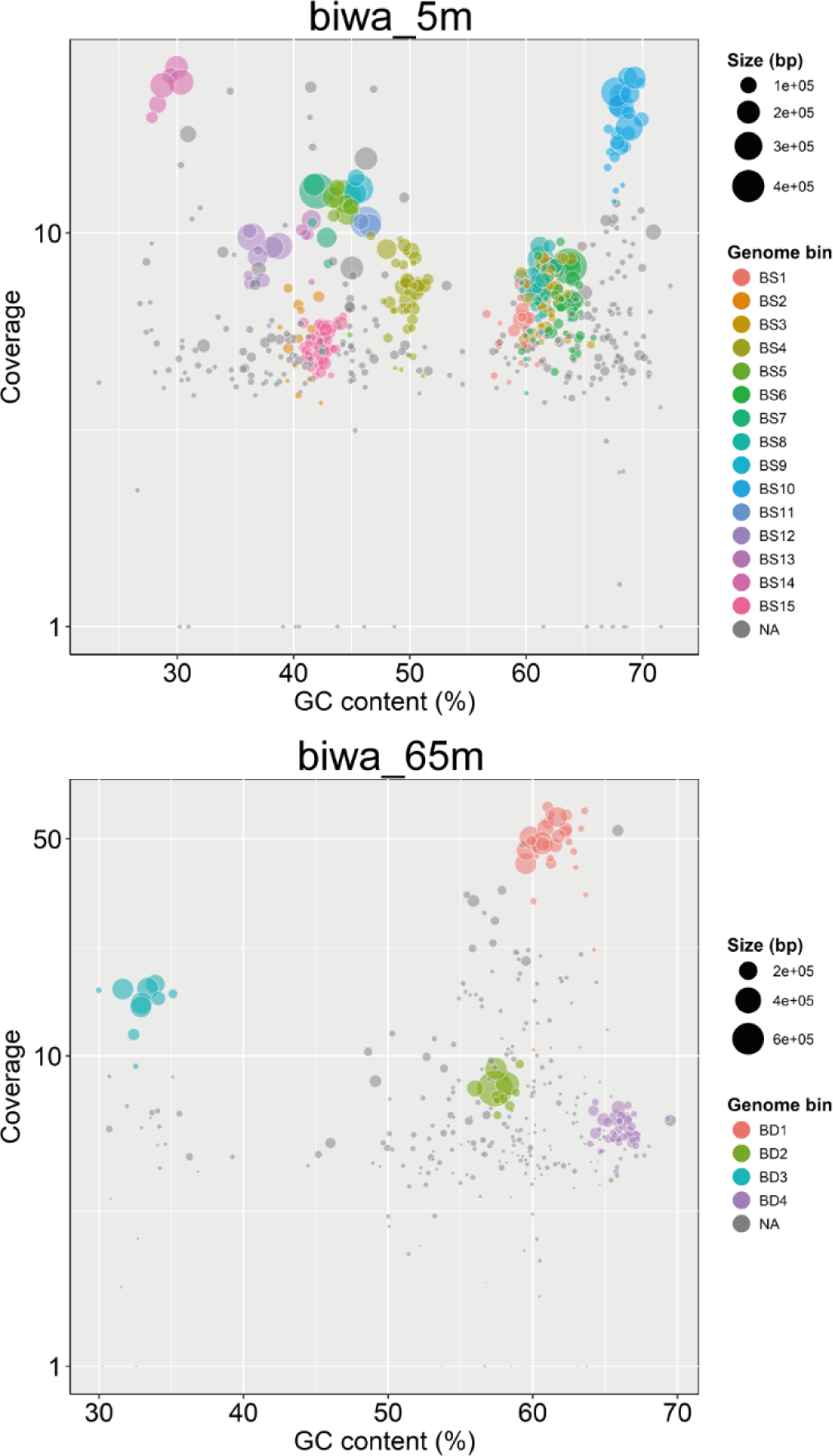
Genome binning of the assembled contigs. Each circle represents a contig, where the color and size represent its assigned bin and total sequence length, respectively. Contigs not assigned to any bin are indicated in gray (named ‘NA’). The x-axis and y-axis represent GC% and genome coverage, respectively.

The nineteen genome bins belonged to seven phyla (Table 2 and Fig. S5). Among these genome bins, ten contained 16S rRNA genes, and many of them showed top hits to uncultured clades; thus, our CCS-based approach was estimated to have truly targeted multiple uncultured prokaryotes. Seven genome bins were predicted to belong to the phylum Actinobacteria, including *Candidatus* Planktophila (BS7), one of the most dominant bacterioplankton lineages in freshwater systems^60,61^. Metagenomic bins affiliated with other dominant freshwater lineages were also recovered, including *Candidatus* Methylopumilus (BS12)^62^, the freshwater lineage (LD12) of Pelagibacterales (BS14)^63,64^, and Nitrospirae (BD2) and *Candidatus* Nitrosoarchaeum (BD3), the predominant nitrifying bacteria and archaea in the hypolimnion, respectively^54,55^. Four bins were affiliated with the phylum Verrucomicrobia (BS6, BS8, BS10, and BD4), in line with a previous study^65^. The BS3 and BD1 genome bins likely represent members of the CL500-11 group (class Anaerolineae) of the Chloroflexi phylum, where BD1 presented the highest coverage of >45×. This group is a dominant group in the hypolimnion of Lake Biwa and is frequently found in deep oligotrophic freshwater environments worldwide^66^. Overall, the phylogeny of the reconstructed genomes likely reflects the major dominant lineages present in the water of Lake Biwa.

### Metaepigenomic analysis

A total of 29 methylated motifs were detected in ten genome bins (Table 3). Their methylation ratios ranged from 19–99%, which can be affected by modification detection power, *i.e.*, these ratios are likely lower than the true methylation levels. Three motifs from the BS12 genome bin contained overlapping sequences (HCAG<u>**C**</u>TKC, BGMAG<u>**C**</u>TGD, and GMAG<u>**C**</u>TKC, where B: G/T/C, D: G/A/T, H: A/C/T, K: G/T, and M: A/C, where the underlined bold face indicates methylation sites) that were likely due to incomplete detection of a single methylated motif or heterogeneous motif sequences between closely related lineages contained within that genome bin. A palindromic motif and five complementary motif pairs that likely reflect double-strand methylation were observed in the BS15 bin (*e.g.*, a pair of <u>**A**</u>GCNNNNNNCAT and <u>**A**</u>TGNNNNNNGCT). It may also be notable that three genome bins from the Chloroflexi phylum (BS1, BS3, and BD1) shared the same motif sequence set (G<u>**A**</u>NTC, TTA<u>**A**</u>, and G<u>**C**</u>WGC, where W: A/T), likely due to evolutionarily shared methylation systems.

**Table 3.**
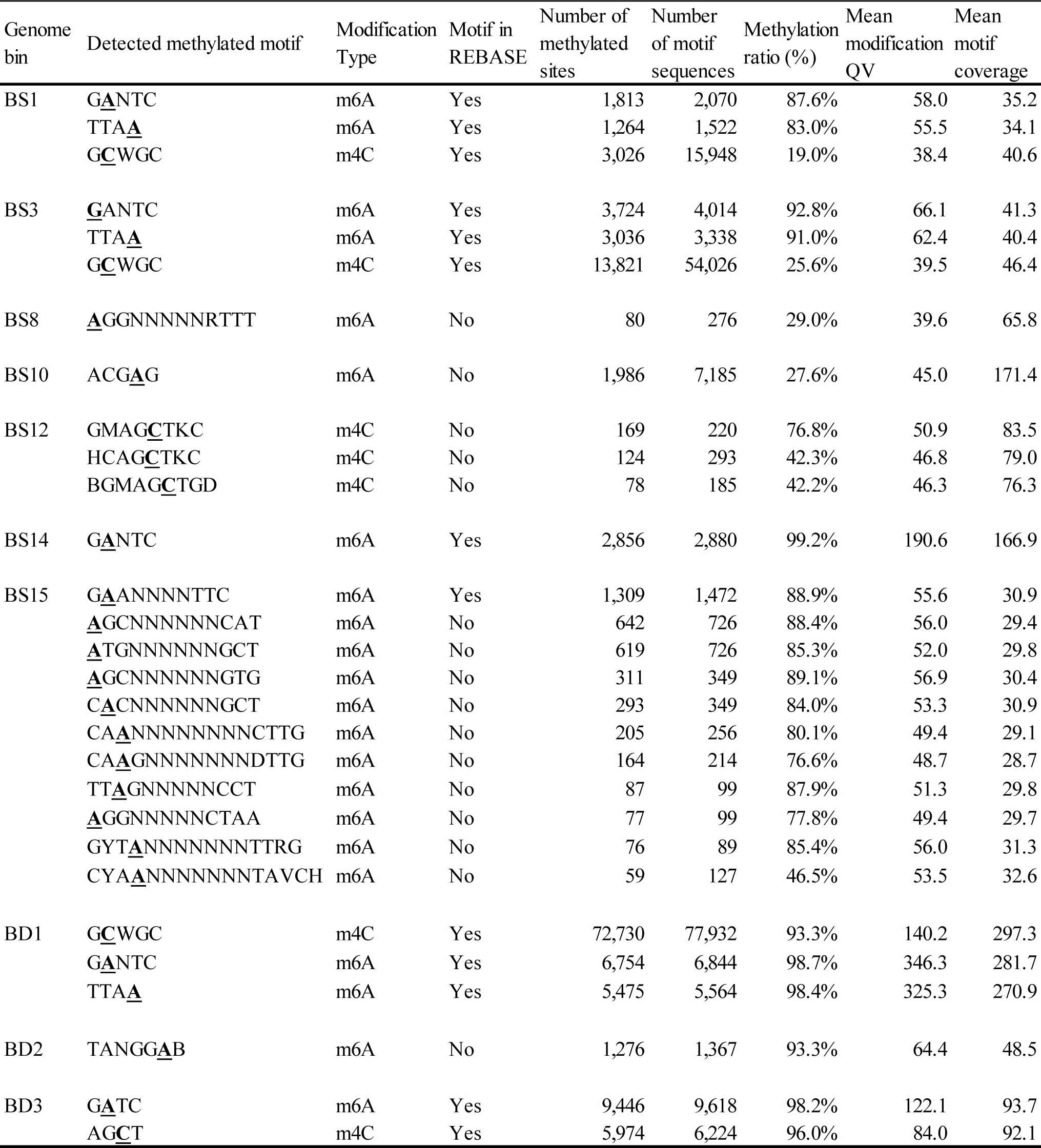
Detected methylated motifs.

Overall, even if such overlapping, complementary, and shared motif sequences are considered, at least 14 motifs still presented no match to existing recognition sequences in the REBASE repository. This result demonstrates the existence of unexplored diversity of DNA methylation systems in environmental prokaryotes, which include many uncultured strains.

### Known MTases that correspond to detected methylated motifs

To identify MTases that can catalyze the methylation reactions of the detected methylated motifs, systematic annotation of MTase genes was performed. Sequence similarity searches against known genes identified 20 MTase genes in nine genome bins (sequence identities ranged from 23–71%) (Table 4). The most abundant group was Type II MTases, followed by Type I and Type III MTases, a trend that is consistent with the general MTase distribution^13,67^. Several genes encoding REases and DNA sequence-recognition proteins were also detected (Table 4). The known motifs of seven of the 20 MTases were matched to those identified in our metaepigenomic analysis (Table 3). For example, the genome bin BD3 contained two MTases, whose recognition motif sequences were AG<u>**C**</u>T and G<u>**A**</u>TC according to the sequence homology-based prediction, which were perfectly congruent with the two motifs detected in our metaepigenomic analysis. It may be notable that these two motifs were also reported in an enrichment-culture study of the closely related genus *Candidatus* Nitrosomarinus catalina^68^ and are therefore likely evolutionarily conserved within their group. In the BS14 bin, a similar one-to-one perfect match was also observed. The two Chloroflexi genome bins BS3 and BD1 were characterized by the same set of three methylated motifs, each of which contained three MTases. No MTase gene was found in the other Chloroflexi bin BS1, likely due to its low estimated genome completeness of 31% (Table 2). Among these MTases, two were predicted to show methylation specificities that were congruent with two of the detected motifs, G<u>**A**</u>NTC and TTA<u>**A**</u> (the other MTase and motif will be discussed in the next section). Collectively, these observations suggest that metaepigenomic analysis is an effective tool for identifying the methylation systems of environmental prokaryotes.

**Table 4.**
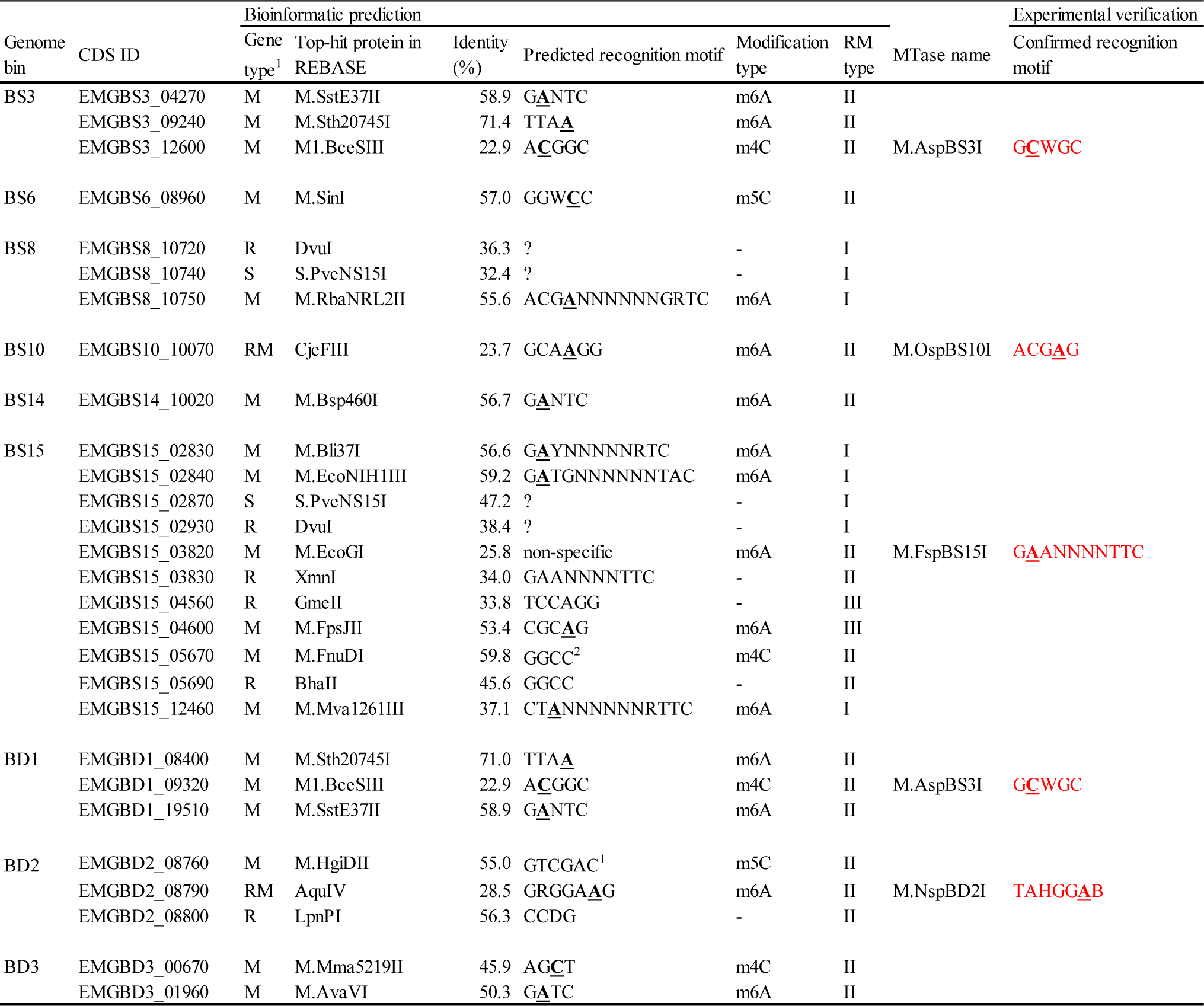
Detected MTases, REases, and specificity subunit genes.

### Unexplored diversity of prokaryotic methylation systems

Among the 20 detected MTases, 13 MTases did not present known recognition motifs that matched those identified in our metaepigenomic analysis (Tables 3 and 4). Although homology search-based MTase identification and recognition motif estimation are frequently conducted in genomic and metagenomic studies, this result suggests that these approaches are not sufficient, and direct observation of DNA methylation is needed to reveal the methylation systems of diverse environmental prokaryotes.

As noted earlier, each of the BS3 and BD1 bins had three MTase genes, two of which were congruent to two of the detected motifs. The other MTase from each bin (EMGBS3_12600 and EMGBD1_09320 in BS3 and BD1, respectively) showed the highest sequence similarity to an MTase that was reported to recognize A<u>**C**</u>GGC; however, the other methylated motif detected in the BS3 and BD1 bins was G<u>**C**</u>WGC.

In the BS15 genome bin, six MTases and eleven methylated motifs were detected, but none of the MTases and motifs matched each other. At the methylation type level, five MTases and all of the methylated motifs were of the m6A type. We predicted that the EMGBS15_03820 MTase, which is estimated to exhibit non-specific m6A methylation activity, is actually a sequence-specific enzyme that recognizes a G<u>**A**</u>ANNNNTTC motif that was detected through metaepigenomic analysis, because the adjacent gene EMGBS15_03830 encodes an REase that targets the same GAANNNNTTC sequence.

In the BS8 genome bin, one MTase and one methylated motif were detected; however, the estimated motif of this MTase was incongruent with the detected motif (the estimated and detected motifs were ACG<u>**A**</u>NNNNNNGRTC and <u>**A**</u>GGNNNNNRTTT, respectively, where R: G/A). This MTase is predicted to function in an RM system because of the existence of the neighboring REase and DNA-sequence recognition protein genes.

In the BS10 genome bin, one MTase and one methylated motif were detected, and their motifs were also incongruent (GCA<u>**A**</u>GG and ACG<u>**A**</u>G, respectively).

In the BD2 genome bin, two MTases and one methylated motif were detected. The two MTases were predicted to display m6A and m5C methylation activities, while the detected motif contained an m6A site. Thus, the former MTase was predicted to catalyze the methylation reaction, although their motifs were again incongruent (GRGGA<u>**A**</u>G and TANGG<u>**A**</u>B, respectively). It should also be noted that these MTases appear to constitute a recently proposed system known as the Defense Island System Associated with Restriction-Modification (DISARM), which is a phage-infection defense system composed of MTase, helicase, phospholipase D, and DUF1998 genes^69^. To our knowledge, this is the first DISARM system identified in the phylum Nitrospirae.

In the BS6 genome bin, one MTase gene was found, but we could not detect any methylated motif, and we therefore anticipate that this MTase gene does not exhibit methylation activity or the corresponding methylation motif was undetected due to the low sensitivity of SMTR sequencing to m5C modification as described previously ^13,14^. However, in the BS12 genome bin, we detected methylated motifs but no MTase genes. We assume that the MTase genes corresponding to this bin were missed due to insufficient genome completeness (although the estimated completeness was 81%), or because these MTase genes have diverged considerably from MTase genes found in cultivable strains, or because thee MTases belong to a new group.

### Experimental verification of MTases with new methylated motifs

Among the MTases whose estimated methylated motifs were not congruent with our metaepigenomic results, we experimentally verified the methylation specificities of the four MTases: EMGBS3_12600 in BS3 (and EMGBD1_09320 in BD1, which has exactly the same amino acid sequence), EMGBS15_03820 in BS15, EMGBS10_10070 in BS10, and EMGBD2_08790 in BD2 (Table 4). We constructed plasmids that each carried one of the artificially synthesized MTase genes, which we then transformed *E. coli* cells that lacked endogenous MTases, forced their expression, and observed the methylation status of the isolated plasmid DNA by REase digestion.

Although the estimated methylated motif sequence of EMGBS3_12600 was A<u>**C**</u>GGC, the unaccounted-for motif sequence observed in BS3 was G<u>**C**</u>WGC. Thus, we hypothesized that the true recognition sequence of EMGBS3_12600 is G<u>**C**</u>WGC. The REase digestion assay showed that TseI (GCWGC specificity) did not cleave the plasmids when EMGBS3_12600 was expressed in the cells, which clearly supports our hypothesis (Fig. 3a). Furthermore, we confirmed that BceAI (ACGGC specificity) cleaved plasmids regardless of whether EMGBS3_12600 was expressed, indicating that the EMGBS3_12600 protein does not show ACGGC sequence specificity (Fig. 3a). Accordingly, we named this protein M.AspBS3I, as a novel MTase that possesses G<u>**C**</u>WGC specificity (Table 4).

**Figure 3.**
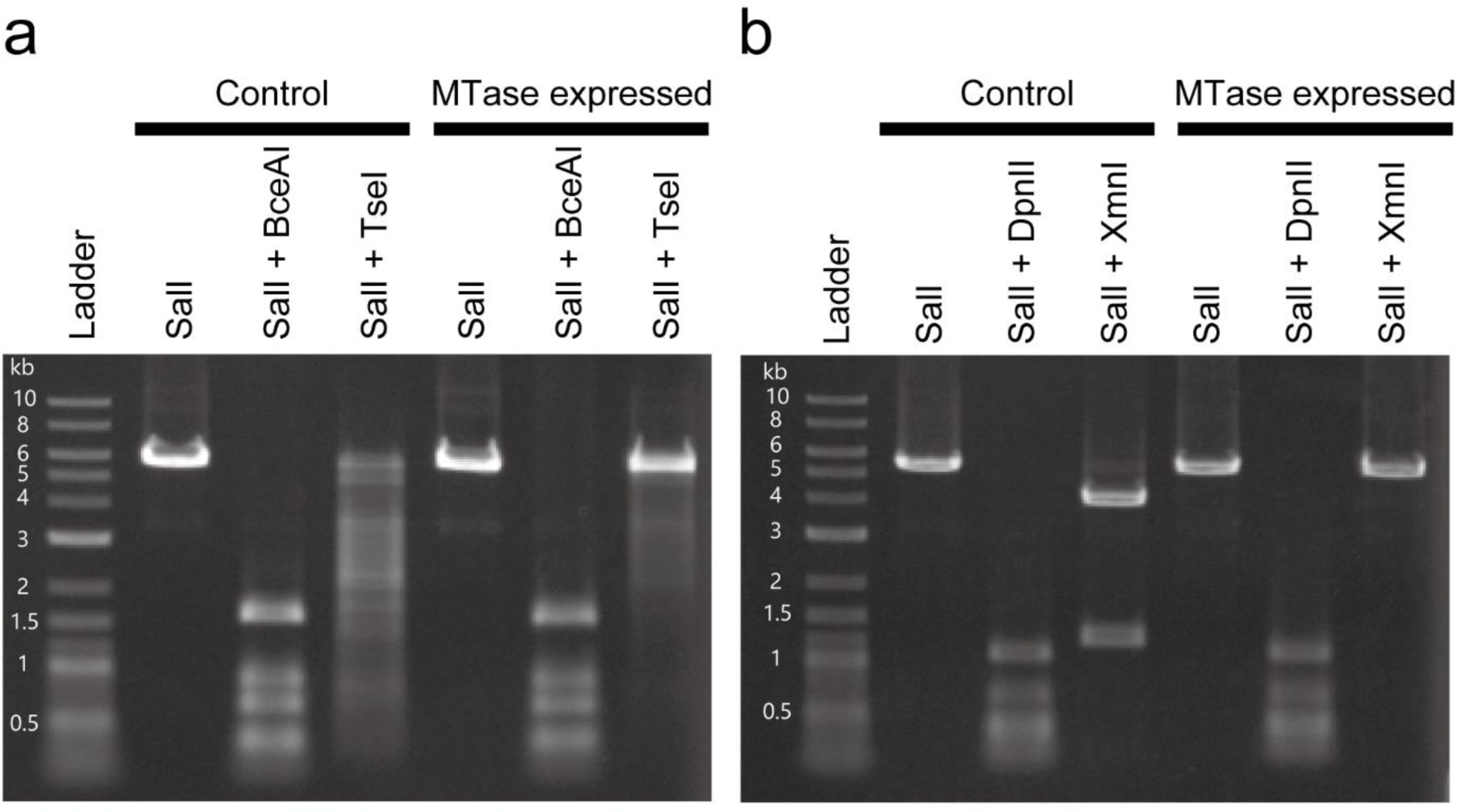
REase digestion assays. **a** Assay of the EMGBS3_12600 gene (and EMGBD1_09320, which has the same amino-acid sequence). BceAI and TseI were used, where the plasmid contained 12 (ACGGC) and 21 (GCWGC) target sites, respectively. Plasmid DNAs were linearized using SalI before the assay. An NEB 2-log DNA ladder was employed as a size marker. **b** Assay of the EMGBS15_03820 gene. DpnII and XmnI were used, where the plasmid contained 27 (GATC) and two (GAANNNNTTC) target sites, respectively.

While the homology-based analysis predicted EMGBS15_03820 as a non-sequence specific MTase, its adjacency to an REase and the results of the metaepigenomic analysis suggested that this MTase presents G<u>**A**</u>ANNNNTTC sequence specificity. The REase digestion assay showed that XmnI (GAANNNNTTC specificity) did not cleave the plasmids only when EMGBS15_03820 was expressed in the cells, which also supports our hypothesis (Fig. 3b). Furthermore, we confirmed that DpnII (GATC specificity) cleaved the plasmids regardless of whether EMGBS15_03820 was expressed, indicating that EMGBS15_03820 is not a nonspecific MTase. We named this protein M.FspBS15I, as a novel MTase that possesses G<u>**A**</u>ANNNNTTC methylation specificity (Table 4).

For EMGBS10_10070 in BS10 and EMGBD2_08790 in BD2, we also conducted REase digestion assays to confirm the recognition motif sequences. Based on the results of the metaepigenomic analysis, their motifs were predicted to be ACG<u>**A**</u>G and TANGG<u>**A**</u>B, respectively. Expression of each gene altered the electrophoresis patterns of the digested plasmids to contain fragments that resulted from inhibition of REase cleavage at the estimated methylation sites (Fig. S6). Furthermore, we additionally conducted SMRT sequencing analysis using the PacBio RSII platform to examine the methylation status of the chromosomal DNA of the *E. coli* transformed with each of the two MTase genes. The results were basically consistent (Table S5): ACG<u>**A**</u>G was actually detected as the methylated motif in *E. coli* transformed with EMGBS10_10070, and we named the protein M.OspBS10I. In the case of EMGBD2_08790, the detected TAHGG<u>**A**</u>B motif was almost the same, but a subset of the estimated TANGG<u>**A**</u>B motif (*i.e.*, TAGGG<u>**A**</u>B was excluded), and this difference could be due to *E. coli*-specific conditions (*e.g.*, cofactors and sequence biases), insufficient data, or inaccuracy of the methylated motif detection method. Regardless of this minor difference, we concluded that EMGBD2_08790 is a novel MTase gene responsible for methylation of the TAHGG<u>**A**</u>B motif and we named the protein M.NspBD2I accordingly.

### Genome bins that lack methylation systems and phage infection

Among the nineteen genome bins, no methylated motifs were detected in nine genome bins (MTase genes were also not detected, except in the BS6 genome bin). This high ratio of methylation-lacking organisms contrasts remarkably with a previous report in which prokaryotic genomes were found to rarely lack DNA methylation systems (<7%)^13^. Notably, those nine genome bins contained seven Actinobacteria bins, indicating that the dominant Actinobacteria in Lake Biwa lack methylation systems, although a number of methylated motifs and corresponding MTases have been reported in Actinobacteria^13^.

Because DNA methylation is known to play a role in opposing phage infection^2–4^, we conducted *in silico* prophage detection to evaluate whether prokaryotes in Lake Biwa tend to be infected by phages. Within the nineteen genome bins, more than one prophage was found in ten genome bins (Table 2 and S6). Among these ten bins, six overlapped with the nine genome bins in which no methylated motifs were identified. The prophages showed little sequence similarity to each other except for two pairs and likely resulted from independent and repetitive infections (Fig. S7). Thus, phage infection and prophage integration appear to frequently occur in prokaryotes that lack DNA methylation systems. We also investigated the presence of CRISPR/Cas systems as another major prokaryotic mechanism against phage infections^70–73^. We identified possible CRISPR arrays in three genome bins, BS3, BS8, and BD3, which exhibit methylation systems but no prophages, although the first two genome bins contained no associated *Cas* genes.

Based on these results, we assume that the possession of prophages is tolerable in lake freshwater environments, and thus, the evolutionary pressure to develop or retain methylation systems is low. These results also suggest that uncultured and cultivable strains may be under different selection pressures regarding DNA methylation systems, and the true diversity of microbial methylation systems must be examined in the future using metaepigenomic approaches.

## Conclusion

The present study demonstrated the effectiveness of the metaepigenomic approach powered by SMRT sequencing and CCS, showing obvious advantages over sequence similarity-based and culture-based methylation system analyses and short-read metagenomics. The CCS reads facilitated metagenomic assembly, binning, and protein sequence-based taxonomic assignment from an environmental sample that contained dominant uncultured prokaryotes. Most importantly, this approach revealed several methylated motifs, including novel ones in environmental prokaryotes, and subsequent experiments identified four MTases responsible for those reactions. The anti-correlation pattern between the presence of prophages and methylation was consistent with past observations that methylation systems inhibit phage infection and phage-mediated genetic exchange, although the underlying ecological background and mechanisms must be examined in the future.

The current throughput of SMRT sequencing may be still insufficient to apply the metaepigenomic approach to more diverse and complex samples. Because deep sequencing coverage (>25× subreads for each DNA strand) is required for the reliable detection of DNA methylation, it is still difficult to obtain sufficient sequencing reads to recover long contigs and detect methylated motifs for ‘rare’ species (typically those with <1% relative abundance). In addition to rapid and ongoing technological advances in SMRT sequencing, the emergence of Oxford Nanopore Technology may provide as another long-read, single-molecule, and methylation-detectable technology^74,75^. Another problem is that the detectable types of DNA modifications are limited (*i.e.*, m4C, m5C, and m6A) with the currently available SMRT sequencing technology, while many other DNA chemical modifications occur in nature^76^. In addition to advances in sequencing methods, novel bioinformatic tools will be critical for metaepigenomic analyses of environmental prokaryotes.

A recent study showed that sets of methylated motifs and MTases can vary widely, even between closely related strains^77^, where metaepigenomics is expected to enable differential methylation analyses between populations. In addition, genus-level conservation of MTases that are not associated with REases is sometimes observed, which suggests that MTases play unexplored adaptive roles, in addition to their functions in combating phages^13,78^. Novel MTases may be adopted for biotechnological uses, such as DNA recombination and methylation analyses^79^. It is envisioned that metaepigenomics of environmental prokaryotes under different sampling conditions and environments will significantly deepen our understanding of the enigmatic evolution of prokaryotic methylation systems and broaden their application potential.

## Author Contributions

SH conceived the study, performed the bioinformatics analyses and experiments, and wrote the manuscript. YO and SN performed the water sampling. AM performed the experiments. AT performed the genomic and metagenomic sequencing. WI conceived the study, wrote the manuscript, and supervised the project. All authors read and approved the final manuscript.

## Acknowledgments

The SMRT sequencing was supported by National Institute of Genetic, Research organization of information and systems, Mishima, Japan. We thank Yoshinori Nii, Masashi Yoshino, and Satoko Fukuda for their helpful suggestions and experimental supports. We are grateful to Yukiko Goda and Tetsuji Akatsuka for their assistance in the field sampling. We also thank Metabologenomics, Inc. for financial support.

